# Mechanical loading and hyperosmolarity as a daily resetting cue for skeletal circadian clocks

**DOI:** 10.1101/2021.10.01.462769

**Authors:** Michal Dudek, Dharshika Pathiranage, Cátia F. Gonçalves, Craig Lawless, Dong Wang, Zhuojing Luo, Liu Yang, Farshid Guilak, Judith Hoyland, Qing-Jun Meng

## Abstract

Daily rhythms in mammalian behaviour and physiology are generated by a multi-oscillator circadian system entrained through environmental cues (e.g. light). Presence of niche-dependent physiological time cues has been proposed, allowing local tissues flexibility of phase adjustment. However, to date, such stimuli have remained elusive. Here we show that cycles of mechanical loading and osmotic stimuli within physiological range drive rhythmic expression of clock genes and reset clock phase and amplitude in cartilage and intervertebral disc tissues. Hyperosmolarity (not hypo-osmolarity) resets clocks in young and ageing skeletal tissues through mTORC2-AKT-GSK3β pathway, leading to genome-wide induction of rhythmic genes. These results advocate diurnal patterns of mechanical loading and consequent daily surges in osmolarity as a bona fide tissue niche-specific time cue to maintain skeletal circadian rhythms in sync.

**One-Sentence Summary:** Circadian clocks in aneural skeletal tissues sense the passage of time through rhythmic patterns of loading and osmolarity

## Main Text

The daily patterns of rhythmic environment on Earth, including light/darkness, temperature fluctuations and availability of food, have profound impact on the physiology and behavior of living organisms. To anticipate and cope with changing demands between day and night, most organisms have evolved an internal cell-intrinsic timing mechanism, the circadian clock. Close alignment of internal circadian rhythms with external environment (known as entrainment) is of paramount importance for health and survival (*1*). In mammals, the current model proposes that the central pacemaker Suprachiasmatic Nuclei (SCN) in the hypothalamus temporally coordinates and adjusts peripheral clocks in all major body organs (*2–4*). Light as a primary and potent entrainment factor resets the central clock which in turn signals directly through neuronal connections or indirectly through hormonal cues to other parts of the brain and the body (*5, 6*). However, one argument for the evolution of local tissue clocks (rather than simply a top down control from the SCN) is that it allows flexibility - peripheral clocks can adopt a different phase relationship with one another (and with the SCN) if circumstances require. If this argument is true, we would expect local clocks to be synchronised to the most physiologically relevant stimuli. One piece of evidence supporting this is the uncoupling of circadian clocks in the liver and other tissues from the SCN upon restricted feeding (*7–11*).

The mammalian skeletal system, especially the articular cartilage in joints and intervertebral disc (IVDs) in the spine, provides a unique model to resolve this question. Cartilage and IVDs are among the most highly loaded tissues, experiencing a diurnal loading cycle associated with daily rest/activity patterns (*12, 13*). In humans, these tissues experience a prolonged period of compression of 2-3.5 MPa during the activity phase, followed by a period of low-load recovery during resting phase (down to 0.1 MPa). These loading patterns lead to diurnal changes in tissue compressive strain, resulting in corresponding changes in the osmotic environment of the cells that may be altered with osteoarthritis, IVD degeneration or systemic risk factors such as obesity and a sedentary lifestyle (*12–15*). Cartilage and IVDs possess intrinsic circadian clocks that drive rhythmic expression of tissue specific genes and become dampened and out of phase in ageing mice. Genetic disruption of cartilage and IVD clocks in mice results in imbalance of anabolic and catabolic processes which subsequently leads to accelerated tissue ageing and degeneration (*16–19*). The aneural and avascular nature of articular cartilage and IVDs makes them less amenable to conventional systemic time cues conveyed through nerves or hormones (*20*). As such, it remains unknown how these skeletal clocks are entrained under physiological conditions. Indeed, we observed no difference in the potency of mouse serum collected during the day vs. night in their ability to synchronise these tissue clocks (Fig. S1), excluding blood-borne serum factors as an effective clock resetting cue for these tissues (*21*). Given the diurnal loading patterns they experience, a priori should expect these peripheral skeletal clocks to be synchronised by some aspects of mechanical loading. However, until now, this has not been shown.

### Mechanical loading resets the circadian clocks in femoral head cartilage and IVD

We tested the role of mechanical loading using an *ex vivo* tissue explant culture model utilising the PER2::Luc reporter mice which allow real-time monitoring of clock protein PER2 using Lumicycle (*22*). A short loading regime (1 hour of 1 Hz, 0.5 MPa compression) resulted in robust amplitude increase of the circadian rhythm in cartilage which was maintained for at least three more days (Fig. 1A). To delineate the best response window within the 24 hour cycle, we applied the same compression protocol at 4 time points 6 hours apart. Only compression applied at the peak of PER2::Luc resulted in a significant increase of circadian amplitude (p<0.001, Fig. 1B, C), with minimal phase shift (Fig. 1D). Compression at other phases resulted in significant phase delay or advance (p<0.001, Fig. 1D). Compression applied at the trough disrupted circadian rhythm, highlighting the importance of the circadian phase in modulating the clock response to loading (Fig1B). The phase shift of the cartilage circadian rhythm was dependent on the magnitude of the force of compression (Fig. 1E).

**Figure 1.**
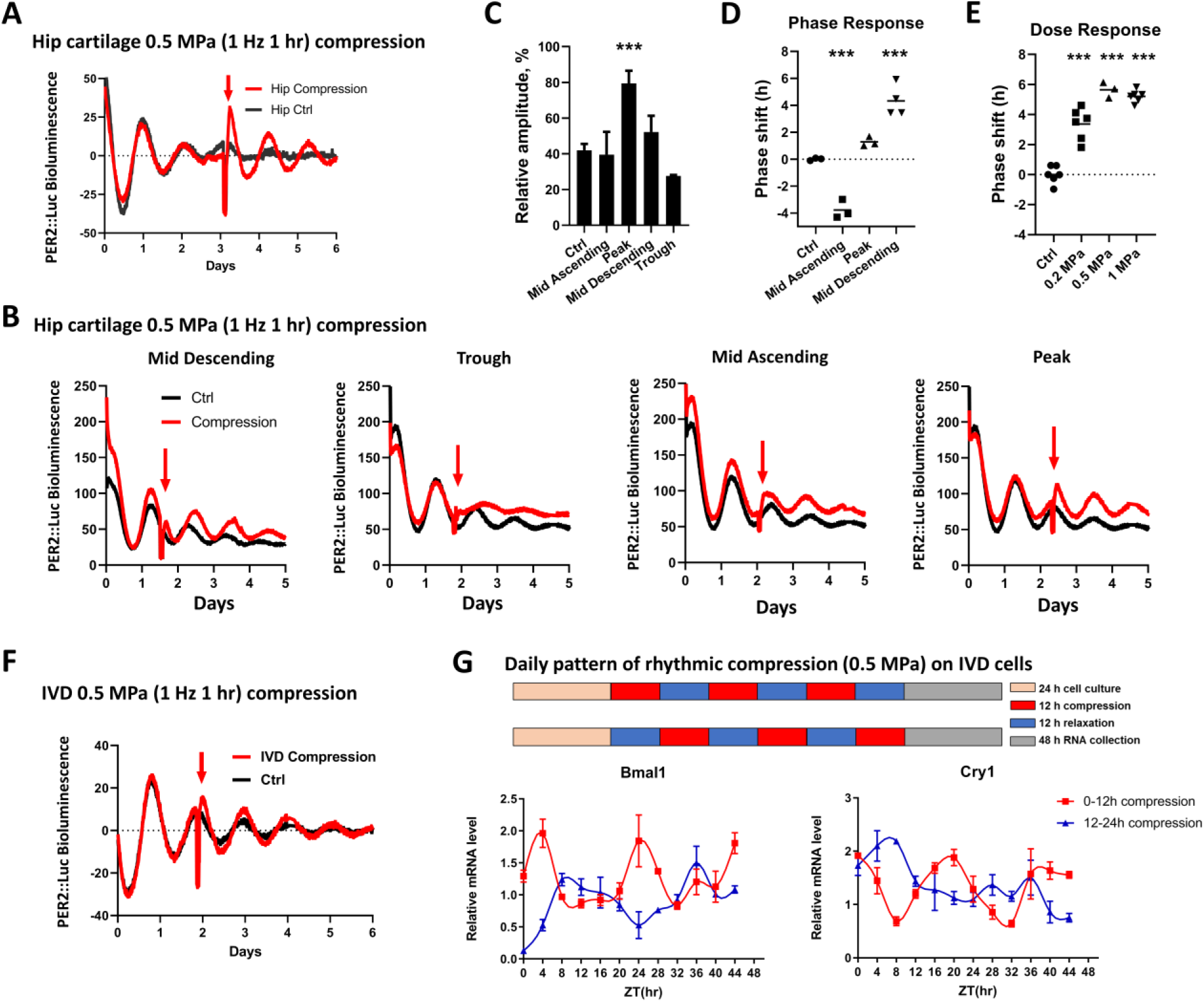
Mechanical loading resets the circadian clock in PER2::Luc cartilage and IVDs in a phase and dose dependent manner. (**A**) Bioluminescence recordings of PER2::Luc femoral head cartilage explants. Red arrow indicates time of compression (0.5 MPa, 1 Hz, 1 hr). Each trace represents the mean of 3 explants. (**B**) Recordings of PER2::Luc cartilage explants subjected to mechanical loading (red arrow) at 6 hr intervals, starting at mid-descending phase. Each trace is the mean of 4 explants. (**C**) Quantification of the PER2::Luc amplitude change in **B**, expressed as % of the amplitude of the peak before loading. One-way ANOVA. ***p<0.001 vs. control. (**D**) Quantification of phase shifts in **B**. ***p<0.001 vs. control. (**E**) Quantification of phase shifts in cartilage exposed to increasing magnitude of compression applied at mid descending phase. ***p<0.001 vs. control. (**F**) Recordings of PER2::Luc IVDs subjected to compression (0.5 MPa, 1 Hz, 1 hr). Each trace is the mean of 3 explants. (**G**) mRNA expression of *Bmal1* and *Cry1* in a rat IVD cell line following 3 cycles of oppositely phased compression (12 hr at 0.5 MPa /12 hr at 0 MPa). Mean ±SD of 3 cultures.

The same loading protocol altered the PER2::Luc rhythm in IVD tissues in a similar manner (Fig.1F and Fig.S2). Next, we tested whether rhythmic loading patterns can entrain circadian rhythms of endogenous clock genes in IVD cells. To this end, rat IVD cells were subjected to oppositely phased mechanical loading cycles of 12-hour loading at 0.5 MPa /12-hour unloading. qPCR showed that the expression of clock genes *Bmal1* and *Cry1* were driven ~180 degrees out of phase (Fig. 1G), further supporting rhythmic loading as an endogenous clock resetting cue for cells in the skeletal system.

### Hyperosmolarity (but not hypo-osmolarity) phenocopies the effect of mechanical loading

The diurnal pattern of mechanical loading of the articular cartilage and IVDs is associated with daily changes in osmotic pressure. Under load, the pressurized interstitial fluid flows to regions of lower pressure, resulting in increased osmolarity. When unloaded the process is reversed, causing a return to normal osmolarity. These fluctuations in osmolarity play an important role in normal skeletal physiology by promoting synthesis of the extracellular matrix and stabilising cellular phenotypes (*23–25*). Therefore, we tested how circadian rhythms in skeletal tissues operate under different baseline osmolarity conditions or when subjected to acute osmotic challenges. PER2::Luc IVD explants were cultured under static osmotic conditions of 230-730 mOsm for 3 days before synchronisation and recording. Here, despite the wide range of osmotic conditions, the intrinsic circadian pacemaking mechanism was still intact with equivalent circadian amplitude and phase, highlighting the ability of these skeletal tissues to adapt to static changes in their environmental osmolarity (Fig.2A). The periodicity was maintained at close to 24 hours despite drastic differences in baseline osmolarity, indicating that circadian period in these skeletal tissues is “osmolarity-compensated” (Fig.2A).

**Figure 2.**
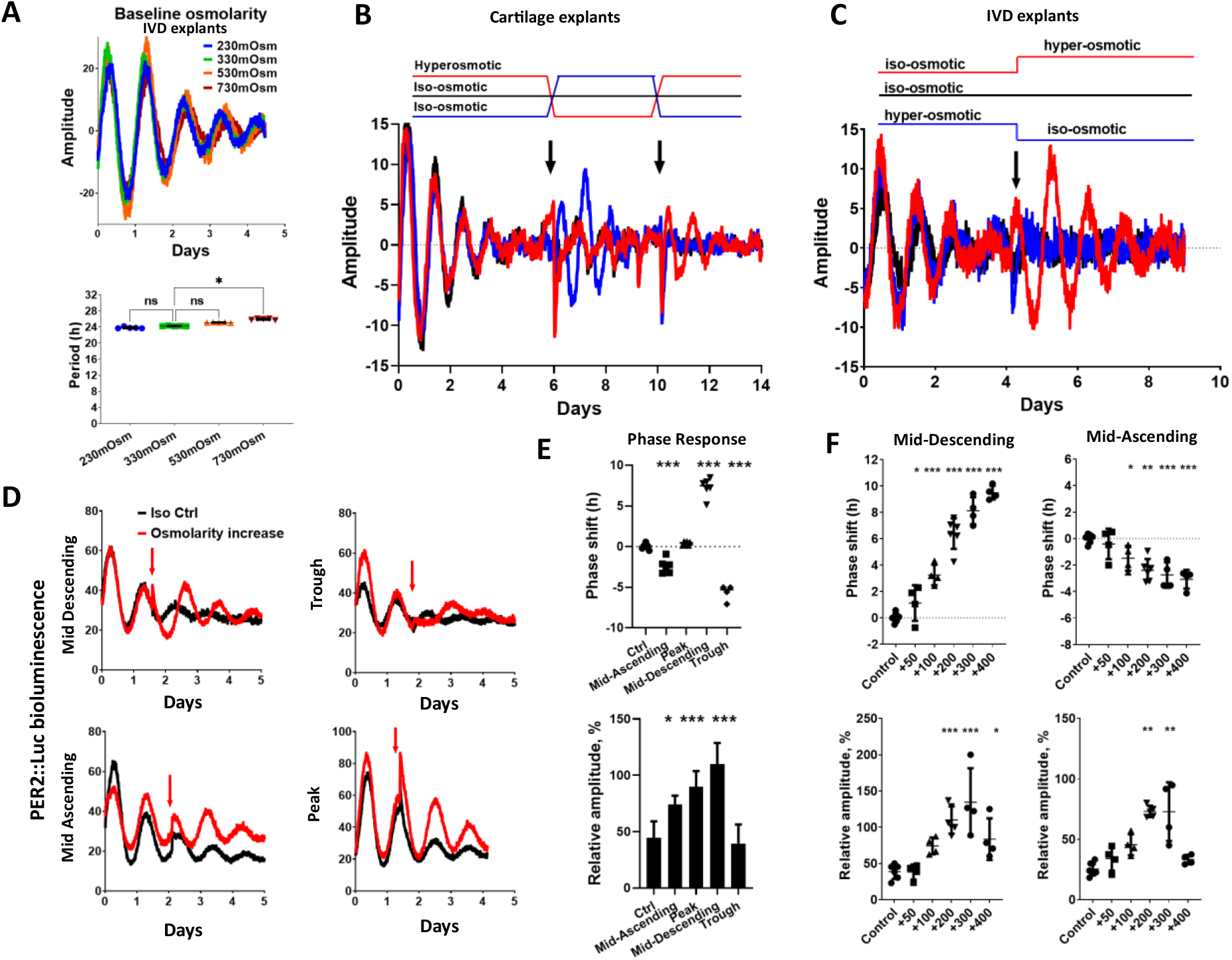
Hyperosmolarity resets the circadian clock in IVDs and cartilage in a phase and dose dependent manner. (**A**) Oscillations and period quantification of PER2::Luc IVD explants cultured in varying baseline osmotic conditions. Mean ±SD, n=5. One-way ANOVA (non-parametric Kruskal-Wallis test). *p<0.05; ns, not significant. (**B**) Oscillations of PER2::Luc cartilage with media changes. Conditioned media in iso- and hyper-osmotic conditions were switched at day 6, then switched back again at day 10. Control cultures were undisturbed. (**C**) PER2::Luc IVD explants cultured in iso- and hyper-osmotic conditions with media swap at day 4. (**D**) Effects of hyperosmolarity (+200 mOsm, applied at 6 hr intervals indicated by red arrow) on PER2::Luc IVD oscillations. (**E**) Quantification of phase shift and amplitude induction in **D.** (**F**) Quantification of dose-dependent phase shift and amplitude induction in IVD explants treated with increasing osmolarity (+50 to +400 mOsm) at mid descending and mid ascending phase of PER2::Luc oscillation. One-way ANOVA. *p<0.05; **p<0.01; ***p<0.001. For panels A-D, each trace represents mean of 3-6 explants.

Next, we analysed how an acute change in osmolarity (as experienced by skeletal tissues on a daily basis) impacts on skeletal clocks. Cartilage tissue explants were placed in iso-osmotic (330 mOsm) or hyperosmotic (530 mOsm) media and allowed to adapt for 3 days before synchronisation and recording. When clocks gradually become desynchronised, the conditioned media were swapped between iso and hyper-adapted explants. The clocks in explants that experienced an increase in osmolarity were resynchronised, with the amplitude recovered to the level at the beginning of the experiment, and robust circadian rhythm continued for the next 4 days. In contrast, explants experiencing a decrease in osmolarity showed gradual loss of rhythmicity, similar to the iso-control explants (Fig.2B). After 4 days the media were swapped back between explants and again the tissues experiencing an increase in osmolarity were resynchronised (Fig. 2B). Clocks in IVD explants showed a very similar response to cartilage, with resynchronisation by increased osmolarity (Fig. 2C and Movie S1).

If altered osmolarity is a key mechanism through which mechanical loading acts on the skeletal clock, it should exert very similar phase and dose dependent effects to mechanical loading. Indeed, a hyperosmotic challenge elicited a very similar direction of phase shifts in a circadian phase-dependent manner comparable to loading (Fig. 2D). The phase response curve (PRC) and the phase transition curve (PTC) demonstrated a type 1 resetting (typical response to a relatively mild physiological clock stimulus) (Fig. S3). The hyperosmolarity induced clock-synchronising effect appears much stronger than loading with improved circadian amplitude at all timepoints except at the trough (Fig. 2E). Similar to loading, the circadian clock response to osmolarity is also “dose” dependent, with a phase delay of up to 9.5 hours (at mid-descending phase) for the +400 mOsm increase (Fig. 2F). Maximal amplitude induction was observed with +300 mOsm condition (Fig. 2F). The dose dependent clock resetting by both mechanical loading and osmolarity was also observed in cartilage and IVD explants from a different clock gene reporter *Cry1*-Luc mouse (*26*) (Fig. S4).

### Daily hyperosmotic challenge synchronises dampened circadian rhythms in both young and ageing skeletal tissues

With the onset of the activity phase mouse cartilage and IVD tissues experience roughly 12 hours of loading within a 24-hour day, leading to increased osmolarity (*20*). Having determined that the acute increase in osmolarity is a clock synchronising factor, we next used a protocol to approximate the diurnal osmotic cycles experienced by skeletal tissues by exposing cartilage and IVD explants to 12 hours hyperosmotic condition and returning them to 12 hours of iso-osmotic medium. A single cycle with a hyperosmotic challenge as low as +100 mOsm synchronised the circadian rhythm in cartilage (Fig. 3A) and IVDs (Fig. 3B) from young mice (2 months old), although, this effect was clearly dependent on the extent of osmotic change and number of cycles applied. When explants were exposed to two daily cycles of +200 mOsm challenges, they showed an even stronger clock amplitude (Fig. 3C, D). In contrast, exposure of tissue explants to a 1 hr hyperosmotic pulse had no significant effect on phase or amplitude (Fig. S5).

**Figure 3.**
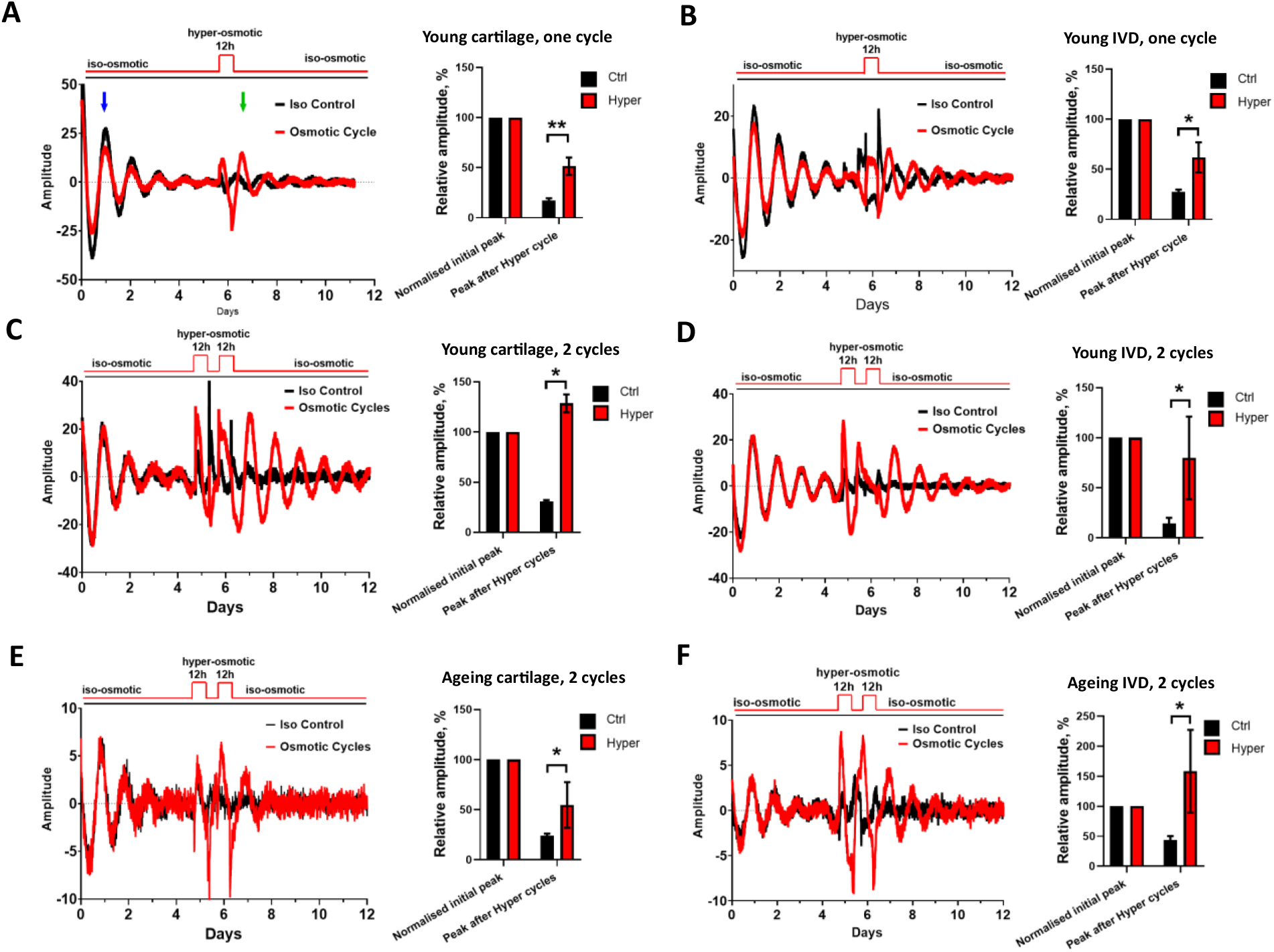
Osmotic cycles synchronise circadian clocks in young and ageing cartilage and IVDs. (**A, B**) Effects of one cycle of osmotic changes on PER2::Luc cartilage explants (A) and IVDs (B) from 2-month old mice. At day 6 explants were exposed to 12 hr of +100 mOsm hyperosmotic medium then returned to iso-osmotic media. Amplitude of the peak after osmotic cycle (green arrow) was quantified as % of the peak 24 hr after start of recording (blue arrow). (**C-F**) Effects of two cycles of hyperosmotic challenges (12 hr of +200 mOsm/12 hr of iso-osmotic media) on cartilage (C, E) and IVDs (D, F) from 2-month old (C and D) or 12-month old (E and F) PER2::Luc mice. Unpaired t-test, *p<0.05; **p<0.01.

We have previously shown that the circadian amplitude of IVD and cartilage rhythms dampen with ageing (*16, 18*). Therefore, we tested whether clocks in ageing skeletal tissues still respond to osmotic cycles. Indeed, exposure of ageing explants to two osmotic cycles resulted in resynchronisation in both articular cartilage (Fig. 3E) and in IVDs (Fig. 3F).

### Hyperosmotic challenge induces rhythmic global gene expression patterns in a cell-autonomous manner

Next, we wanted to investigate how cellular circadian clocks in isolated cells respond to hyperosmotic challenge. Clock-unsynchronised primary chondrocytes from PER2::Luc mice exhibit only a low amplitude oscillation. Exposure to +200 mOsm increase in osmolarity augmented PER2::Luc amplitude (Fig. 4A). Similar observations were made in a human NP115 (IVD) cell line (Fig. S6A). Strikingly, the same hyper-osmotic stimuli disrupted circadian oscillation in U2OS cells, a human osteosarcoma cell line widely used as a cellular model of circadian clocks (Fig. S6B). As such, the osmolarity-entrainment of circadian clock is clearly cell type dependent and indicative of cellular adaptation to their local niche.

**Figure 4.**
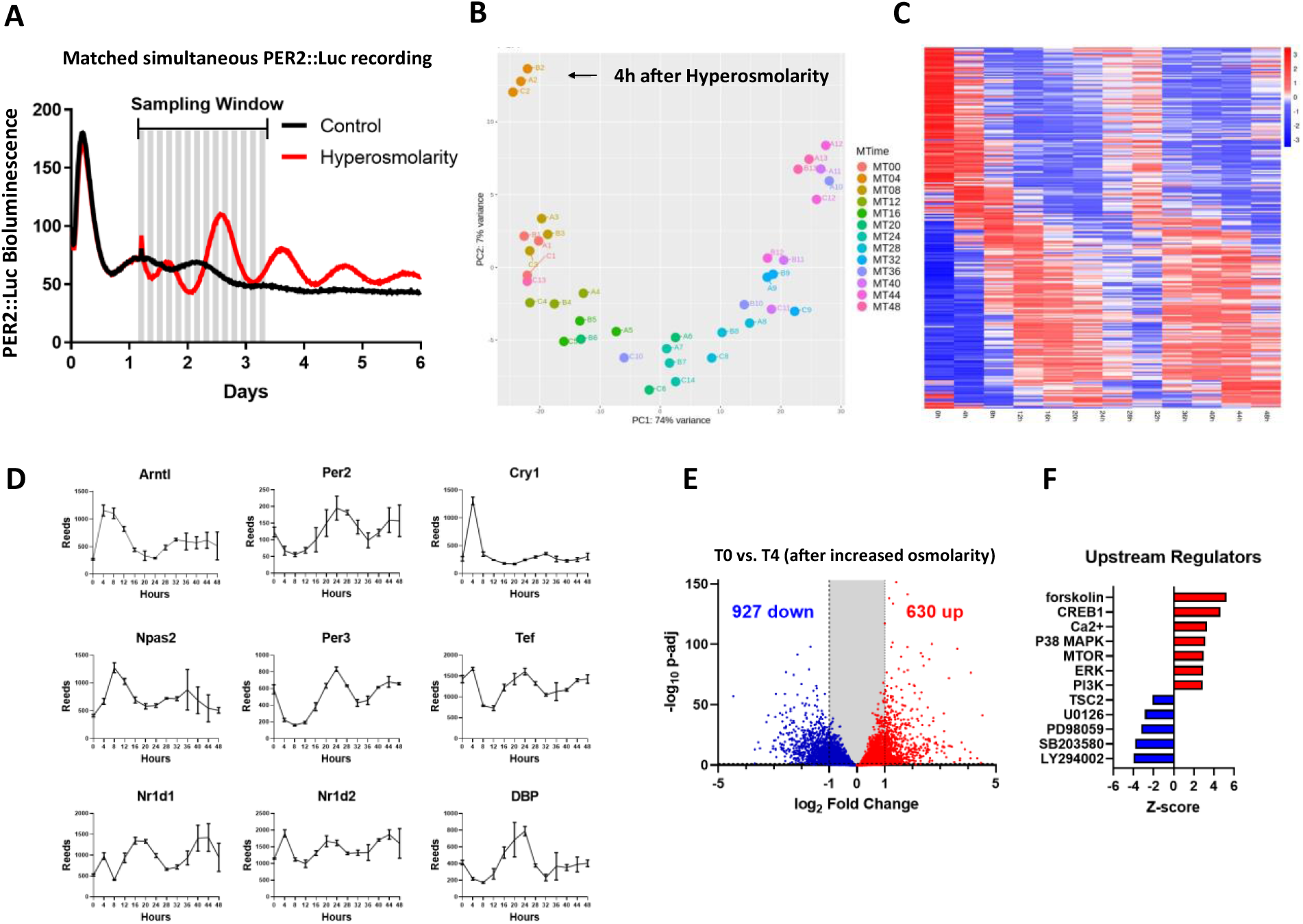
Hyperosmolarity induces wide scale rhythmic gene expression at the transcriptome level. (**A**) Bioluminescence recording of primary chondrocytes isolated from PER2::Luc mice. Cells were not synchronised at the beginning of the experiment. After 30 hr, media osmolarity was increased by +200 mOsm. Parallel samples were harvested every 4 hr for RNA isolation and RNAseq analysis. (**B**) Principal component analysis showing correlation of RNAseq replicates and a developing trend over time. (**C**) Heatmap showing gene expression patterns of 254 rhythmic genes (BHQ<0.05) following increase of osmolarity. (**D**) RNAseq raw data for core circadian clock genes following increase of osmolarity (Mean ±SD, N=3). (**E**) Volcano plot showing differentially expressed genes between T0 and T4 (4 hr after increase in osmolarity). 1,557 genes showed significant changes (>2-fold, p-adj<0.05). (**F**) IPA prediction of top upstream regulators for differentially expressed genes between T0 vs. T4. Red, positive correlation; blue, negative correlation.

We next determined to what extent hyperosmolarity can synchronise circadian rhythms of gene expression at the transcriptome level by circadian time-series RNA sequencing in primary mouse chondrocytes. The PER2::Luc reporter allowed us to track clock rhythms in parallel cultures (Fig. 4A). mRNA samples were collected every 4 hours for 2 full circadian cycles, starting with samples just before the osmotic increase (Time 0h). Principal component analysis of the RNAseq results revealed the circadian time of sampling as the biggest factor separating the samples (Fig. 4B). MetaCycle analysis revealed 1,312 rhythmic genes using integrated P<0.05 cut-off threshold (254 rhythmic genes with a BHQ<0.05 cut-off), including most core clock genes (Fig. 4C, D; Data S1, S2). IPA analysis revealed significantly rhythmic pathways including “Circadian Rhythm Signalling”, “Protein Ubiquitination Pathway”, “mTOR Signalling” and “Autophagy” (Data S3).

### Hyperosmolarity- induced clock resetting involves ERK1/2 and p38, but not CREB or Ca^2+^

Following hyperosmolarity, 1,557 genes showed significant differential expression (with >2-fold change) between T0 and T4 (Fig. 4E, Data S4). Upstream Regulators analysis based on differential genes revealed Forskolin (a known clock synchronising compound), CREB1, p38, Ca^2+^, ERK and mTOR as most significant candidates (Fig. 4F, Data S5). Treatment with forskolin had a synchronising effect on IVD clocks (Fig.S7A). However, inhibitors of CREB had no effect on blocking the hyperosmolarity induced rhythms (Fig. S7B, C). Equally, hyperosmolarity (or mechanical loading) elicited similar clock changes in IVD, cartilage and primary chondrocytes regardless of the presence or absence of calcium in the medium, nor the presence of Gd^3+^, a wide-spectrum cell membrane calcium channel blocker (Fig. S8A-D). Western blotting confirmed activation of ERK1/2 and p38 upon hyperosmotic challenge in mouse primary chondrocytes (Fig.S9A, C), consistent with earlier reports (*27, 28*). Inhibition of ERK1/2, p38, or MEK (upstream of ERK1/2) blocked the amplitude effect of hyperosmotic challenge in PER2::Luc IVDs (Fig. S9B, D; Fig.S10A, B). However, these inhibitors did not have any significant effect on loading-induced clock resetting (Fig. S11A, B).

### mTORC2-AKT-GSK3β as a convergent pathway for loading and hyperosmolarity induced clock resetting

Another pathway that featured prominently in Upstream Regulators analysis of the upregulated genes by hyperosmolarity was mTOR (Fig. 4F, Data S5). Therefore, we tested two mTOR inhibitors, Torin1 (blocking both mTORC1 and mTORC2 complex) and Rapamycin (primarily blocking mTORC1). While Torin1 completely blocked the increase in amplitude following hyperosmotic challenge in cartilage and IVD explants, Rapamycin had no effect (Fig. 5A, B, Fig. S11), indicating involvement of mTORC2. AKT is a known phosphorylation target of mTORC2 (*29*) and can be activated by hyperosmolarity in renal cells (*30*). Indeed, pre-treatment of cartilage explants with an AKT inhibitor prevented the increase in circadian amplitude following hyperosmotic challenge (Fig. 5C). AKT is a known upstream regulator of GSK3β activity, a key clock regulating kinase implicated in the regulation of the stability and/or nuclear translocation of PER2, CRY2, CLOCK, REV-ERBα and BMAL1 (*31, 32*). Indeed, hyperosmotic challenge increased GSK3β phosphorylation at Ser9 and Ser389 (corresponding to reduced activity) (Fig. 5D). Concordantly, inhibition of GSK3β activity by lithium had a synchronising effect on IVD clocks similar to that of hyperosmolarity (Fig. 5E). Pre-treatment with Torin1 (but not Rapamycin) or an AKT inhibitor decreased GSK3β phosphorylation at Ser9 and Ser389 following hyperosmotic challenge (Fig. 5F-H). We next verified whether the above mechanisms were also involved in clock resetting by mechanical loading. Pre-treatment with the mTORC2 or AKT inhibitor (but not Rapamycin) completely blocked the mechanical loading induced increase in circadian clock amplitude (Fig. 5I, J, Fig. S12). Supporting our findings, mechanical strain in mesenchymal stem cells initiates a signalling cascade that is uniquely dependent on mTORC2 activation and phosphorylation of AKT at Ser-473, an effect sufficient to cause inactivation of GSK3β (*33*). The convergent mTORC2-AKT-GSK3β pathway supports osmolarity change as an important downstream factor in mediating loading-entrained skeletal clocks.

**Figure 5.**
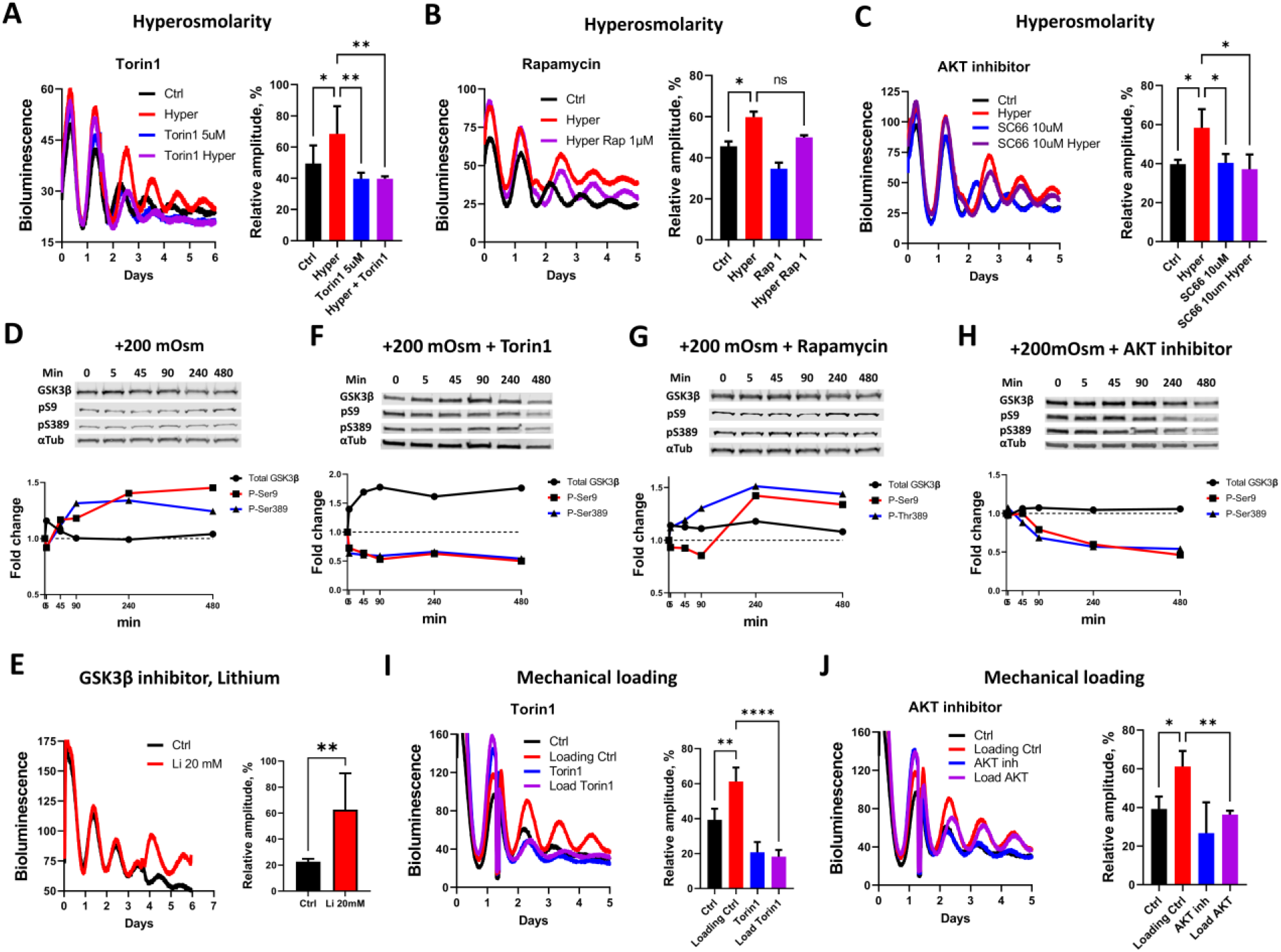
The mTORC2-AKT-GSK3β pathway is a convergent mechanism mediating the loading and hyperosmolarity elicited clock entrainment. **(A-C)** Effects of mTORC1/2 inhibitor Torin1 (A), mTORC1 inhibitor Rapamycin (B) and AKT inhibitor SC66 (C) on blocking the hyperosmolarity (+200 mOsm) -induced clock amplitude in PER2::Luc cartilage explants. **(D)** WB showing total and phosphorylation levels of GSK3β at Ser9 and Ser389 in response to +200 mOsm change in osmolarity in mouse primary chondrocytes. **(E)** Bioluminescence recording and amplitude quantification of PER2::Luc IVD explants treated with 20 mM lithium (an inhibitor of GSK3β). **(F-H)** Effects of Torin1 (F), Rapamycin (G) and AKT inhibitor SC66 (H) on blocking the hyperosmolarity-induced phosphorylation of GSK3 β at Ser9 and Ser379 in mouse primary chondrocytes. **(I, J)** Effects of Torin1 (I) or the AKT inhibitor SC66 (J) on blocking the loading (0.5 MPa, 1 Hz, 1 hr) induced clock amplitude change in PER2::Luc cartilage explants. One-way ANOVA for A, B, C, I and J. *p<0.05; **p<0.01; ****p<0.0001. Unpaired t-test for E. **p<0.01.

Taken together, we have identified daily cycles of mechanical loading and associated rhythmic changes of osmolarity within the physiological range as bona fide entrainment cues for skeletal circadian clocks. These time cues are particularly important for tissues such as the IVD and cartilage because they are isolated from vascular, lymphatic and nervous systems, yet their circadian rhythms are sensitive to age-related dampening (*20*). We show that repeated daily osmotic cycles lead to resynchronisation of circadian rhythm in ageing tissue explants, supporting a hypothesis that reduced rest/activity patterns in ageing individuals could contribute to skeletal clock disruptions *in vivo.*

The clock resetting effect by loading and hyperosmolarity is circadian phase dependent. An osmotic increase or loading at the peak of PER2 (which corresponds to the activity onset in mice) does not shift the phase but strengthens the amplitude of skeletal clocks, which could have implications for the timing of exercise. Regular exercise is thought to be beneficial for the maintenance of skeletal muscle, joint and IVD (*15*, *34*). Indeed, scheduled exercise was shown to reset the circadian clocks in a time-dependent manner in the SCN, skeletal muscle and adrenal gland (*35–37*). However, these studies have largely focused on metabolic and stress related pathways downstream of exercise, instead of the role of mechanical loading or osmolarity.

A study reported a 30 min hypertonic pulse of +600 mOsm led to clock phase shifts in MEFs, NIH3T3 and U2OS cells (*38*). However, +600 mOsm is well outside the physiological range of most human or mouse cells. In the nucleus pulposus region of IVD where extracellular osmolarity is the greatest, osmolarity fluctuates between 400 to 550 mOsm while in cartilage it varies between 350 and 480 mOsm (*20*), depending on the time of day. Indeed, we found chronic +200 mOsm was well tolerated by skeletal cells but showed a severe detrimental effect on clocks in U2OS cells. As such, the clock entrainment effect by physiological fluctuations of osmolarity is clearly specific to the unique skeletal tissue niche.

Based on our findings, it is plausible to predict that daily surges of osmolarity, at the starting phase of activity, may serve as a critical mechanism for skeletal tissues to sense the passage of time and keep its endogenous rhythm in sync with the external environment. As such, our findings resolve the question of how the cartilage and IVD clocks are entrained and confirm tissue-unique inputs as a feature of circadian organisation in mammals. These physiological inputs may allow flexibility for cartilage/IVD clocks to be uncoupled from the SCN clock. Indeed, when typical nocturnal animals are energetically challenged by food shortage or thermal stress, they switch temporal niche from nocturnal to diurnal (*39–40*). These biological insights will also help us understand how ageing and a sedentary lifestyle impact on circadian rhythms and facilitate design of time-prescribed interventions to maintain musculoskeletal health.

## Acknowledgments

We thank J Takahashi (UT Southwestern Medical Center, US) and M Hastings (MRC LMB, Cambridge, UK) for the PER2::Luc and *Cry1-Luc* mouse lines. We thank GC Van Den Akker, TJ Welting and JW Voncken (Maastricht University, Netherlands) for their kind provision of the human NP115 cell line. We thank the Genomics Core Facility (A Hayes and L Zeef) and the Bioimaging Facility (D Spiller) at the University of Manchester for their kind assistance with RNAseq and imaging studies. We thank R Lucas and J Allen for their comments on our manuscript.

## Author contributions

Conceptualization: QJM, JAH, MD

Methodology: MD, DP, CL, DW, LY, FG

Investigation: MD, DP, CG, DW, CL, LY, ZJL, QJM, JAH

Visualization: MD, CG, CL, DW.

Funding acquisition: QJM, JAH, ZJL

Project administration: QJM, JAH

Supervision: QJM, JAH

Writing – original draft: QJM, MD, JAH

Writing – review & editing: LY, ZJL, FG, CG, LY

## Data and materials availability

All data are available in the main text or the supplementary materials.

## Materials and Methods

### Animals

All animal studies were performed in accordance with the 1986 UK Home Office Animal Procedures Act. Approval was provided by the local ethics committee. Mice were maintained at 20-22°C, with standard rodent chow available *ad libitum* and under 12:12 hr light dark schedule (light on at 7 am; light off at 7 pm). The PER2::Luc mice carry the firefly luciferase gene fused in-frame with the 3’ end of the *Per2* gene, creating a fusion protein reporter [1]. *Cry1*-luc mice were previously described [2]. In this transgenic mouse line, the endogenous *Cry1* promoter is fused with a luciferase reporter.

### Tissue explant culture, mechanical loading and osmotic challenge

Mouse articular cartilage tissue explants were prepared as described before [3] by detaching the cartilage from subchondral bone of the femoral head using a scalpel. From the same mice whole IVDs were dissected from the lumbar region of the spine as we described before [4]. Explants were cultured and recorded at 37°C, 5% CO_2_ in DMEM/F12 without phenol red and without serum. Osmolarity of the medium was increased using sorbitol and measured using freezing point osmometer. Mechanical loading of tissue explants was performed in BioPress compression plates on FX-5000 system (FlexCell).

### Bioluminescence recording and imaging

Bioluminescence from PER2::Luc tissue explants or clock reporter cells were recorded in real-time in Lumicycle (Actimetrics) as described before [3, 4]. For recording, 100 μM luciferin was added to the medium. When required, explants were synchronized using 100 nM dexamethasone for 1 hr before bioluminescence recording. Where appropriate traces were normalized using 24 hr moving average. Live tissue bioluminescence of PER2::Luc IVD explant was imaged using a selfcontained Olympus LuminoView LV200 microscope and recorded using a cooled Hamamatsu ImageEM C9100-13 EM-CCD camera [3]. Images were taken every hour for 6 days and combined using ImageJ software (NIH).

### Mouse primary chondrocyte culture and cell lines

Primary chondrocytes from 5 day old mice were isolated according to a published protocol [5]. Briefly, 5 day old C57bl/6 PER2::Luc mice were sacrificed by decapitation. Knee, hip and shoulder joints were dissected, and any soft tissue removed. Joint cartilage was subjected to predigestion with collagenase D (3 mg/mL) in DMEM two times for 30 min at 37°C with intermittent vortex to remove soft tissue leftovers. Subsequently, cartilage was diced using a scalpel and digested overnight at 37°C. Cells were dispersed by pipetting and passed through 70 μM cell strainer. Cell suspension was then centrifuged and the pellet was re-suspended in DMEM/F12 with 10% FBS and plated in T75 flasks. Cells were passaged only once before experiments. Immortalised human nucleus pulposus (NP) cell line NP115 was described previously [6]. The cell line was stably transfected with a pT2A-*mPer2*::Luc reporter construct (a kind gift from Kazuhiro Yagita, Kyoto Prefectural University of Medicine) using nucleofection, followed by single cell colony selection. U2OS cells stably transfected with *Per2*-Luc plasmids were kindly gifted by Patrick Nolan (MRC Harwell).

### Calcium imaging of primary chondrocytes

2 days before calcium imaging, primary chondrocytes were plated onto 4 chamber 35-mm glass-bottomed dishes (Greiner Bio-One). 30 min before imaging medium was changed to 500 μL DMEM/F12 without FBS (with or without 1 mM calcium). Next, 0.5 μL of 1 mM Fluo-4 AM calcium dye (Thermo Fisher) was added to each chamber of cells before incubation on the Zeiss Exciter confocal microscope stage at 37°C in humidified 5% CO2. Imaging was performed using 488 nm excitation wavelength and 520 nm band-pass filter for emission and Fluar 40× NA 1.3 (oil immersion) objective. Image capture was performed with the Zeiss software Aim version 4.2 utilizing the Autofocus macro [7]. Fluorescence analysis was performed on all cells in the field of view using the Zeiss software Aim version 4.2.

### Cyclic mechanical loading on rat NP cell line

Immortalized rat NP cells were kindly generated and donated by Di Chen (Shenzhen Institutes of Advanced Technology) and maintained in DMEM/F-12 (1:1) (Gibco, USA) containing 10% FBS (Invitrogen, USA) and 1% penicillin/streptomycin (Gibco) at 37°C under 5% CO2 and 20% O2 [8]. Compression was conducted 24 hr after cell attachment. Compressive loading was exerted by a pressure incubator according to our previous studies [9, 10]. The rat NP cells were divided into two groups and held under antiphase compression cycles (alternating 12:12 hr cycles of 0.5 MPa/0 MPa for 3 days). After compression cycles, the two groups of NP cells were cultured in conventional incubator and mRNA was extracted every 4 hr for 48 hr before RNA isolation and qRT-PCR. Primers sequence (5’-3’, Rattus norvegicus):

*Bmal1* forward, GACTTCGCCTCCACCTGTTCAA;

*Bmal1* reverse, GCAGCCCTCATTGTCTGGTTCA.

*Cry1* forward, GCGGAAACTGCTCTCAAGGA;

*Cry1* reverse, CCCGCATGCTTTCGTATCAG.

### Circadian time series RNAseq in primary chondrocytes

Mouse primary chondrocytes were passaged into 6-well plates. Upon reaching confluency, media were changed to fresh and ~30 hr later osmolarity was increased by +200 mOsm using sorbitol. T0 sample was collected just before the addition of sorbitol. Then samples were collected every 4 hr for 48 hr. mRNA was extracted using RNeasy micro kit (Qiagen) according to the manufacturer’s protocol. Quality and integrity of the RNA samples were assessed using a 2200 TapeStation (Agilent Technologies). Libraries were generated by the Genomic Technologies Core Facility using the TruSeq® Stranded mRNA assay (Illumina, Inc.) according to the manufacturer’s protocol. Briefly, total RNA (0.1-4 μg) was used as input material from which polyadenylated mRNA was purified using poly-T, oligo-attached, magnetic beads. The mRNA was then fragmented using divalent cations under elevated temperature and then reverse transcribed into first strand cDNA using random primers. Second strand cDNA was then synthesised using DNA Polymerase I and RNase H. Following a single ‘A’ base addition, adapters were ligated to the cDNA fragments, and the products then purified and enriched by PCR to create the final cDNA library. Adapter indices were used to multiplex libraries, which were pooled prior to cluster generation using a cBot instrument. The loaded flow-cell was then paired-end sequenced (76 + 76 cycles, plus indices) on an Illumina HiSeq4000 instrument. Finally, the output data was demultiplexed (allowing one mismatch) and BCL-to-Fastq conversion performed using Illumina’s bcl2fastq software, version 2.20.0.422.

### Bio-informatic analysis

Unmapped paired-end sequences were tested by FastQC (http://www.bioinformatics.babraham.ac.uk/projects/fastqc/). Sequence adapters were removed and reads were quality trimmed using Trimmomatic_0.39 [11]. The reads were mapped against the reference mouse genome (mm10/GRCm38) and counts per gene were calculated using annotation from GENCODE M21 (http://www.gencodegenes.org/) using STAR_2.7.7a [12]. Normalisation, Principal Components Analysis, and differential expression was calculated with DESeq2_1.28.1 [13]. Only genes exceeding 50 counts in at least one timepoint were used subsequently. Analysis of rhythmic genes was done with MetaCycle_1.2.0 [14]. MetaCycle is an algorithm for detection of rhythmicity that combines results of three methods (Lomb-Scargle periodograms, JTK_CYCLE and Arser) that involve least-squares fits to sinusoidal curves, detecting monotonic orderings of data across ordered independent groups and autoregressive spectral estimation. In subsequent analysis we used the Meta_2d function of MetaCycle which provides integrated p value of the three methods as well as BHQ adjusted p value. For Ingenuity Pathway Analysis (Qiagen) we used the integrated p value list to investigate rhythmic pathways. Differentially expressed genes between T0 and T4 were used for Upstream Regulator analysis with threshold set at adjusted p<0.05 and -Log_2_ Fold Change ≤1. RNAseq raw data were deposited to ArrayExpress repository (accession no. E-MTAB-11040).

### Western Blotting

Western blotting was performed according to standard procedures. All primary antibodies were purchased from Cell Signaling Technology ERK1/2 (#4695), pThr202/pTyr204 ERK1/2 (#4370), p38 (#8690), pThr180/pTyr182 p38 (#4511), GSK3β (#12456), pSer9 GSK3β (#5558), except for pSer389 GSK3β (14850-1-AP) which was purchased from Proteintech and αTubulin (T9026) purchased from Sigma Aldrich. Primary antibodies were used in 1:1000 dilution. Secondary antibodies (LI-COR IRDye 800CW and 680RD) were used in 1:20,000 dilution. WB were imaged using the LI-COR Odyssey Imaging System and quantified using Image Studio.

### Reagents

Dexamethasone and Forskolin were purchased from Sigma. Inhibitors 666-15, PD184352, SB203580, Rapamycin, Torin1 and SC66 were purchased from Tocris. KG-501 and GDC-0994 were purchased from Cambridge Bioscience Ltd.

### Statistical analysis

Data were evaluated using Two-tailed Student’s t-test, One way ANOVA or non-parametric, two-tailed Mann-Whitney test. Following ANOVA individual comparisons were performed using Fisher’s LSD test. Results were presented as mean ± SD from at least three independent experiments. Differences were considered significant at the values of *P < 0.05, **P < 0.01 and ***P < 0.001.

---

**Figure S1.**
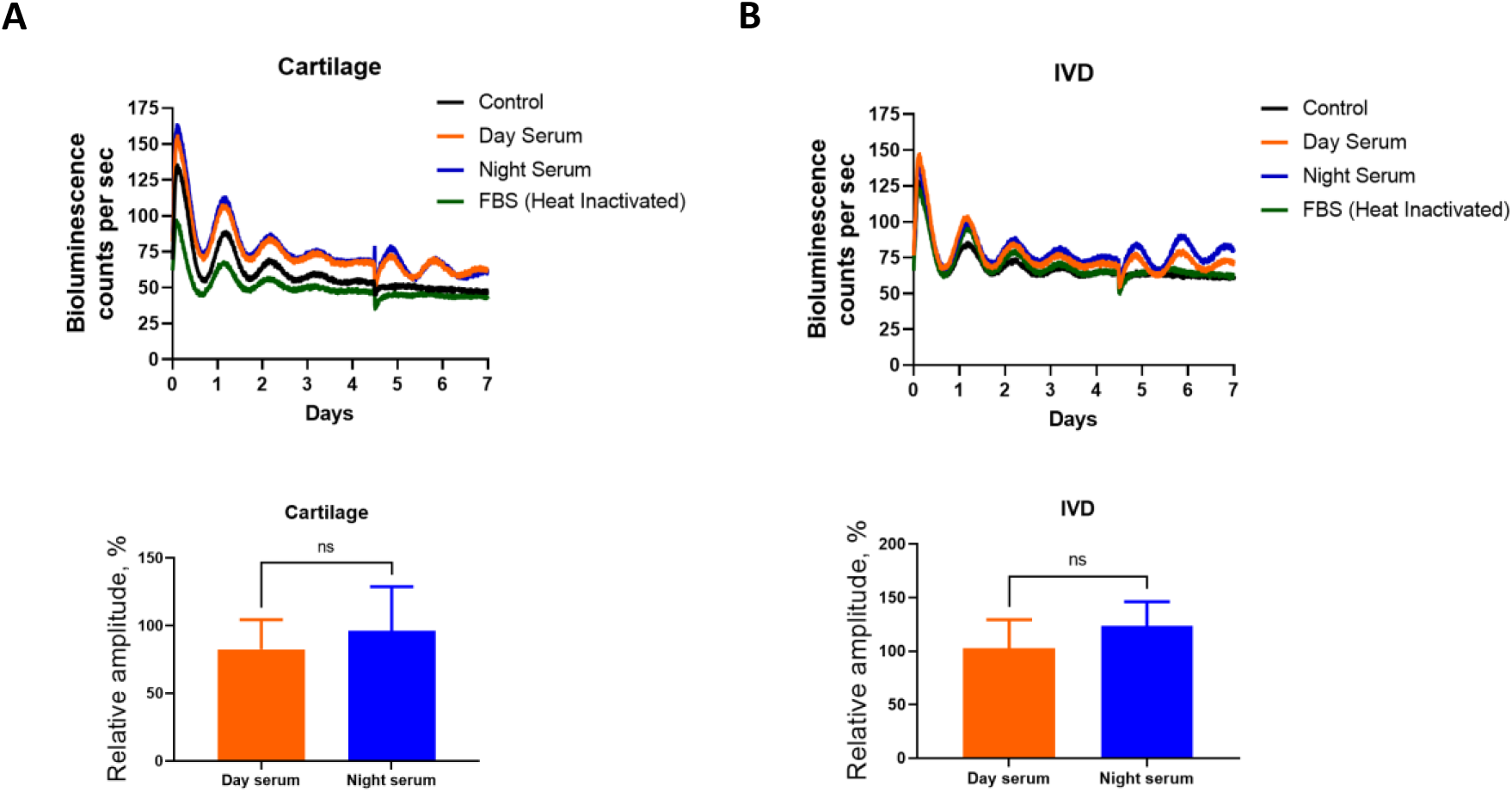
There is no difference between day serum and night serum in their ability to synchronise skeletal clocks. Bioluminescence recordings of PER2::Luc femoral head cartilage (**A**) and IVD (**B**) explant. At day 4, 10% mouse serum harvested during the day (orange) or night (blue) was added to the culture. 10% heat inactivated FBS (green) and 10% volume of fresh medium (black) were included as controls. Amplitude of the peak after treatment was quantified as % of the peak at day 1 of recording. Unpaired t-test; n=3; ns- not significant.

**Figure S2.**
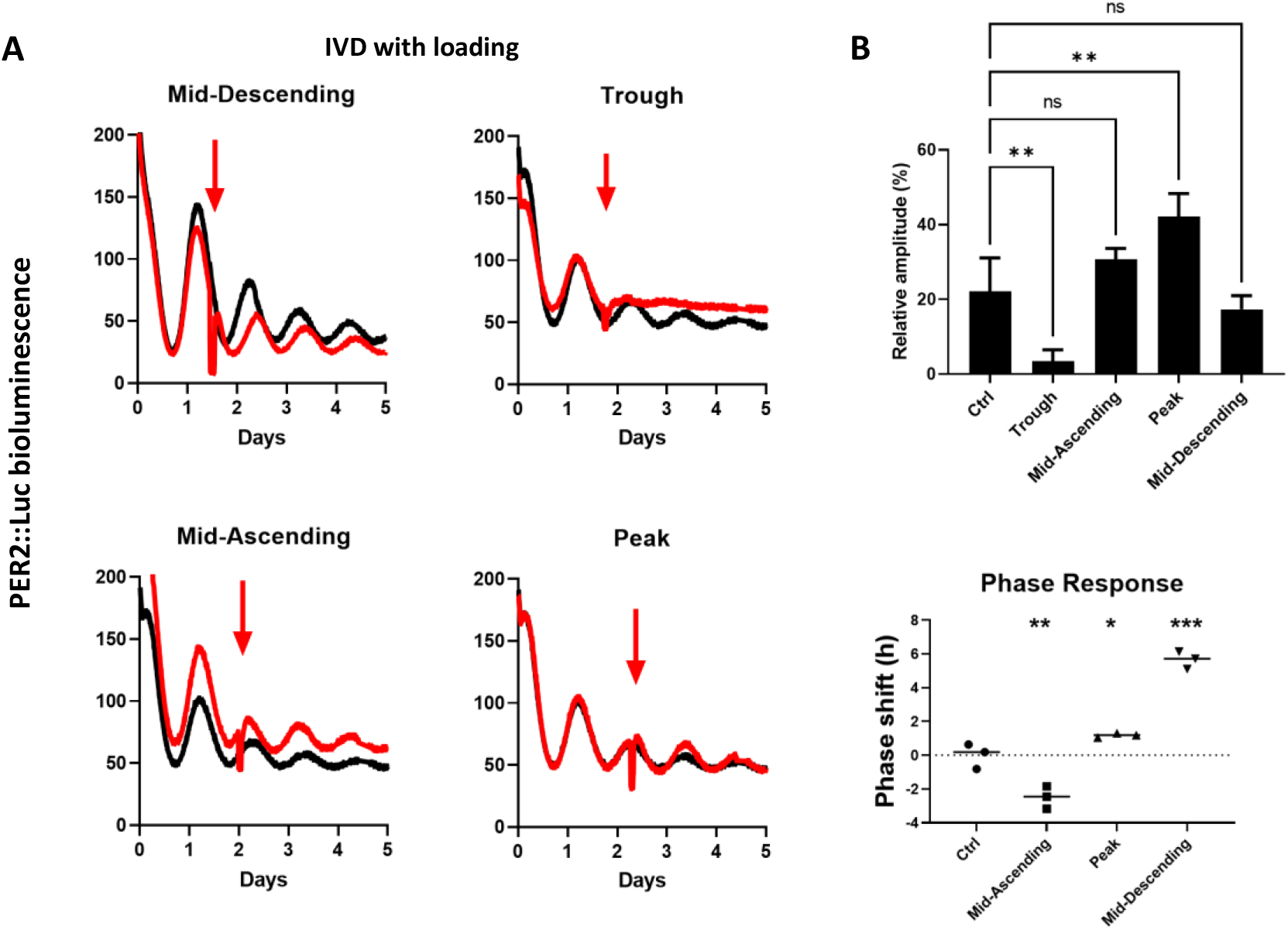
Mechanical loading resets clock phase and amplitude in PER2::Luc IVD explants. (**A**) Bioluminescence recordings of PER2::Luc IVDs subjected to mechanical loading (red arrow) at 6 hr intervals. Each trace represents mean of 3 explants. (**B**) Quantification of the amplitude (top) and phase shift (bottom) after mechanical loading in **A**. Amplitude is expressed as % of the amplitude of the peak before loading. Mean ± SD. One-way ANOVA, *p<0.05; **p<0.01; ***p<0.001.

**Figure S3.**
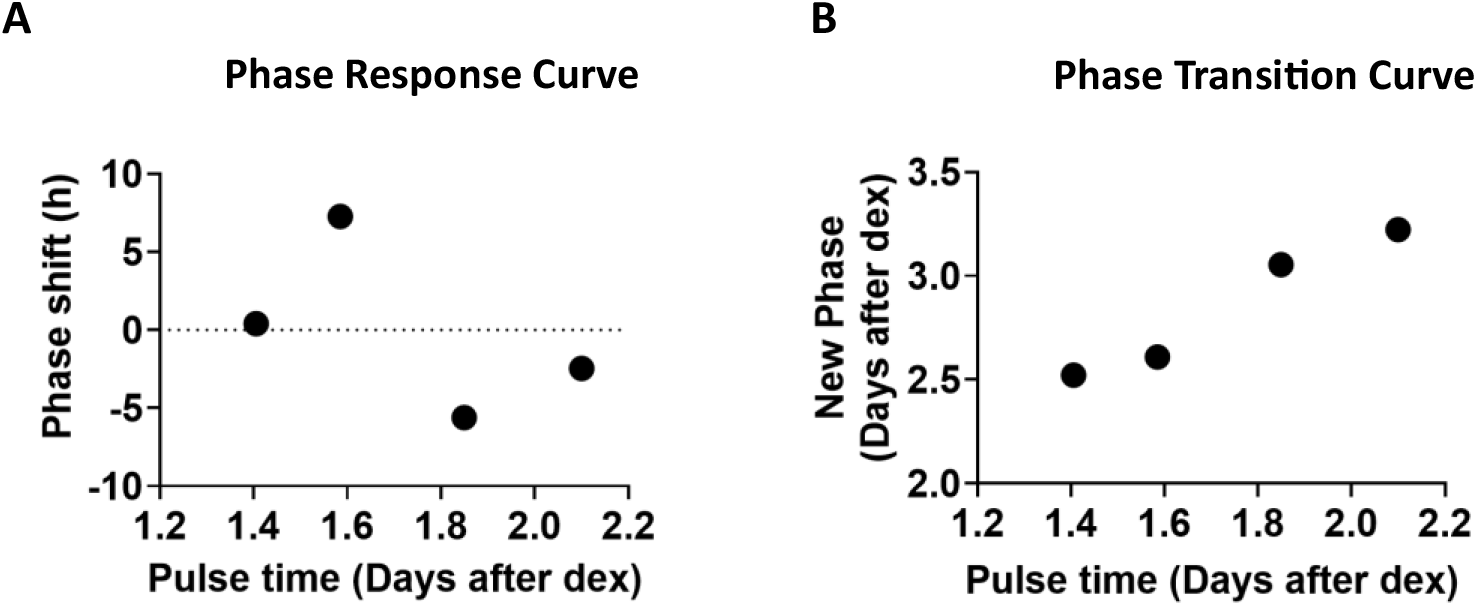
Hyperosmolarity elicits a type 1 resetting of the circadian clock phase in IVD explants. Quantification of the phase response curve (**A**) and phase transition curve (**B**) after exposure of IVDs to +200 mOsm increase in osmolarity in relation to data shown in Figure 2D.

**Figure S4.**
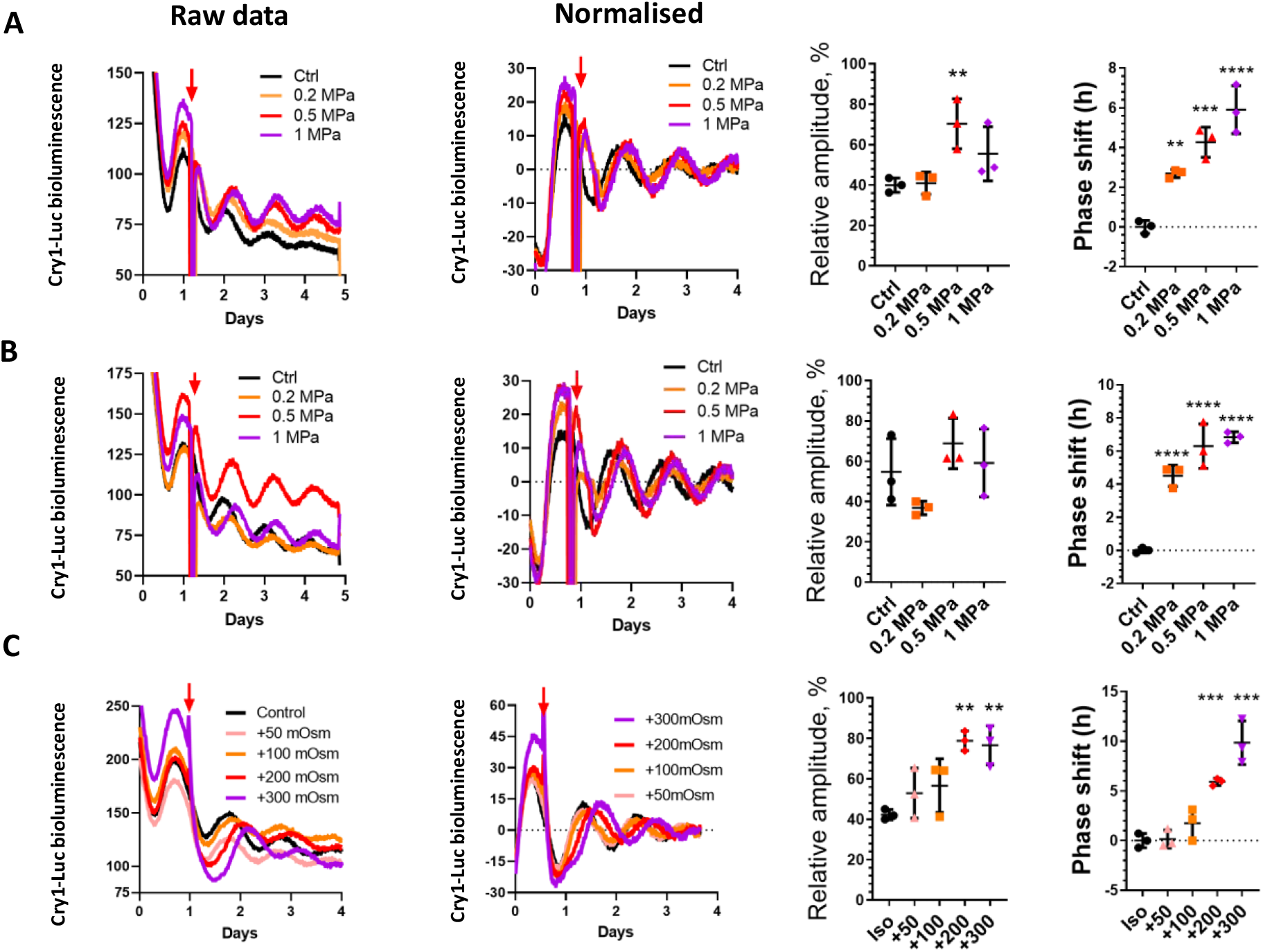
Clock resetting effect by loading and hyperosmolarity in cartilage and IVD explants from *Cry1-Luc* reporter mouse. (**A, B**) At ~30 hr of recording (red arrow) cartilage (A) or IVD (B) explants were subjected to mechanical loading (1 Hz, 1 hr) at compression magnitudes of 0.2 MPa, 0.5 MPa or 1 MPa. Each trace represents mean of 3 explants. (**C**) At ~30 hr of recording (red arrow) IVD explants were exposed to a range of osmolarity increase. Each trace represents mean of 3 explants. Left panels, raw bioluminescence counts; middle panels, normalised; right panels: Quantification of amplitude and phase shift after loading or hyperosmolarity (mean ±SD). One-way ANOVA, **p<0.01; ***p<0.001; ****p<0.0001.

**Figure S5.**
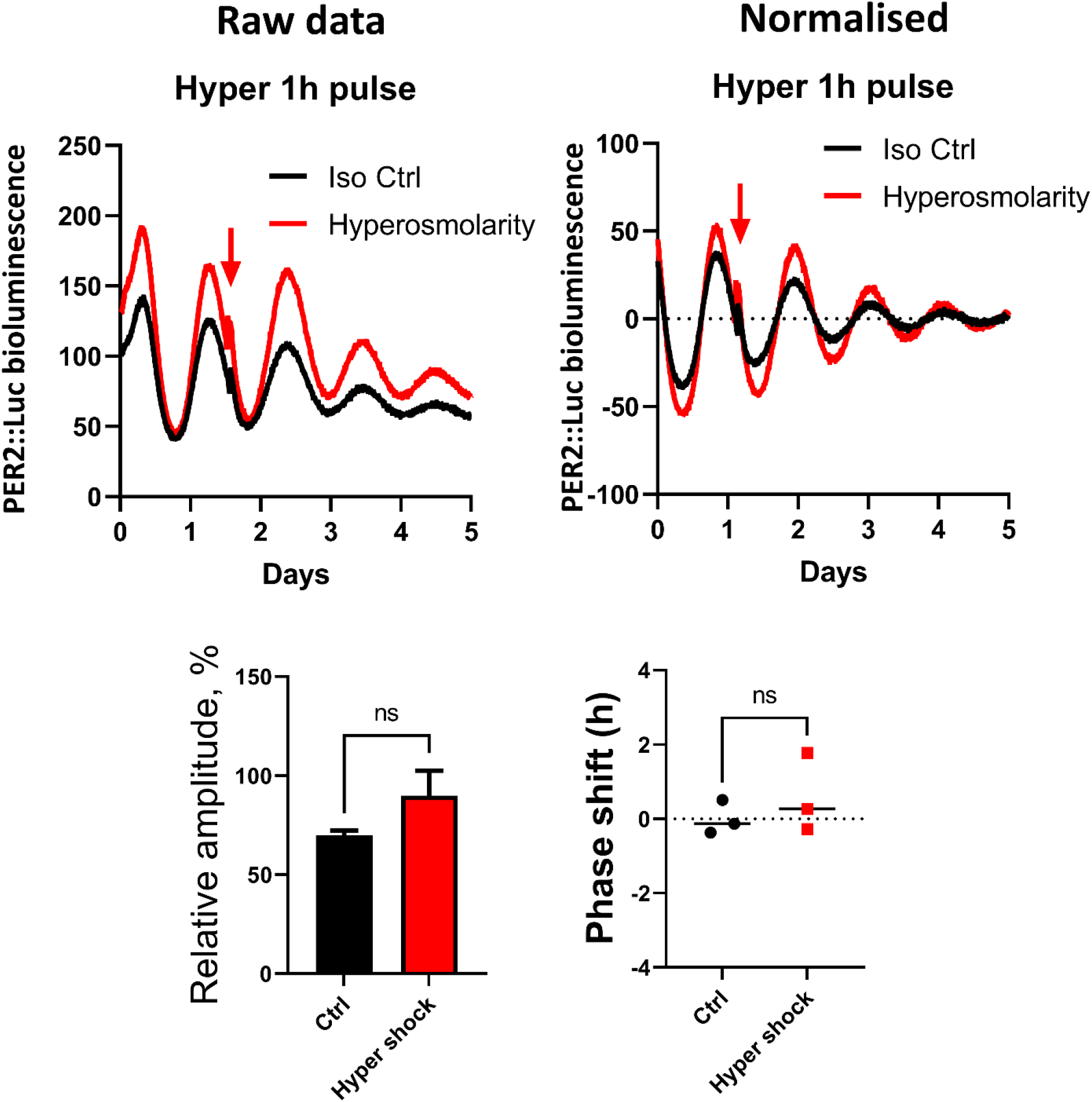
A short pulse (1 hr) of hyperosmolarity is not sufficient to reset IVD clocks. PER2::Luc IVD explants were exposed to 1 hr +200 mOsm hyperosmolarity pulse (red arrow) and returned to iso-osmotic media. Each trace is mean of 3 explants. Amplitude is expressed as % of the amplitude of the peak before loading. Mean ±SD. Unpaired t-test, ns- not significant.

**Figure S6.**
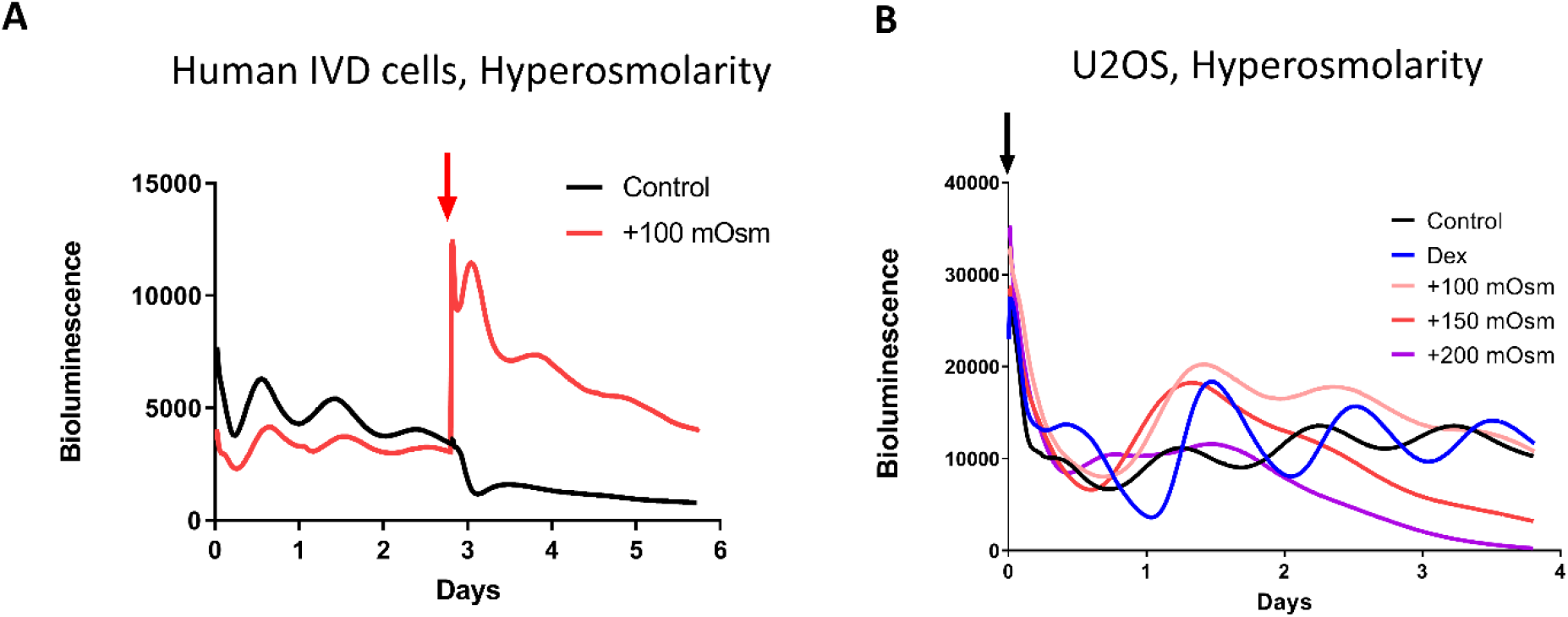
Cell type dependent effect of hyperosmolarity on clocks. (**A**) Bioluminescence recordings of a human IVD cell line NP115 carrying a *Per2*-Luc reporter exposed to +100 mOsm increase (red arrow). (**B**) Bioluminescence recordings of a human U2OS cell line carrying a *Per2-*Luc reporter. At the beginning of the experiment (black arrow) cells were synchronised using dexamethasone (blue) or exposed to an increase in osmolarity. Black trace is the unsynchronised and untreated control.

**Figure S7.**
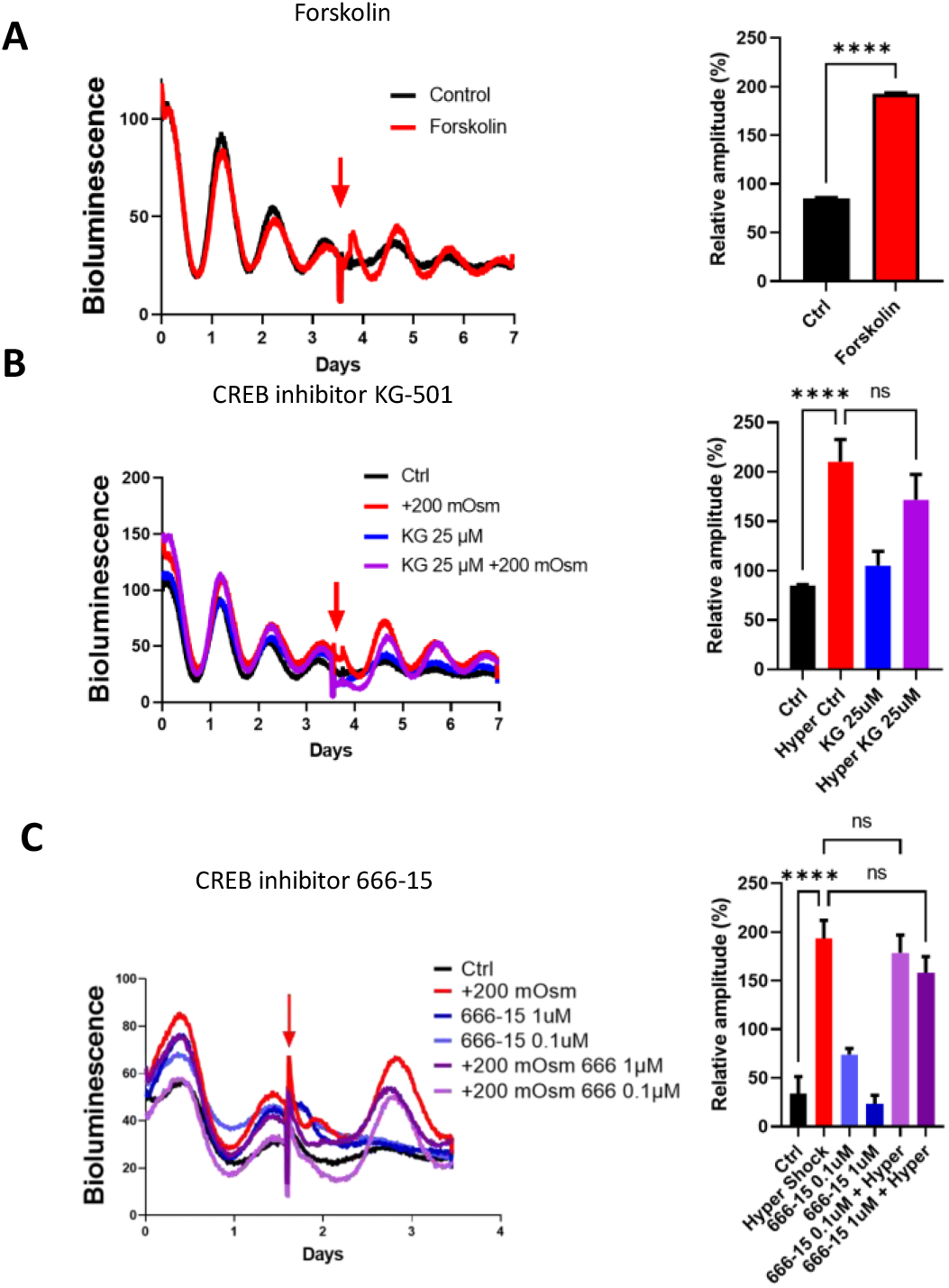
CREB activity is not required for hyperomolarity induced clock resetting in IVDs. (**A**) Bioluminescence recording and amplitude quantification of PER2::Luc IVD explants treated with forskolin (red arrow) ( **B**) Explants were exposed to +200 mOsm increase in osmolarity with or without CREB inhibitor KG-501. (**C**) Explants were exposed to +200 mOsm increase in osmolarity with or without CREB inhibitor 666-15. Each trace represents mean of 3 explants. Amplitude of the peak after treatment is expressed as % of the amplitude of the peak before treatment. Mean ±SD. One-way ANOVA. ****p<0.0001; ns- not significant.

**Figure S8.**
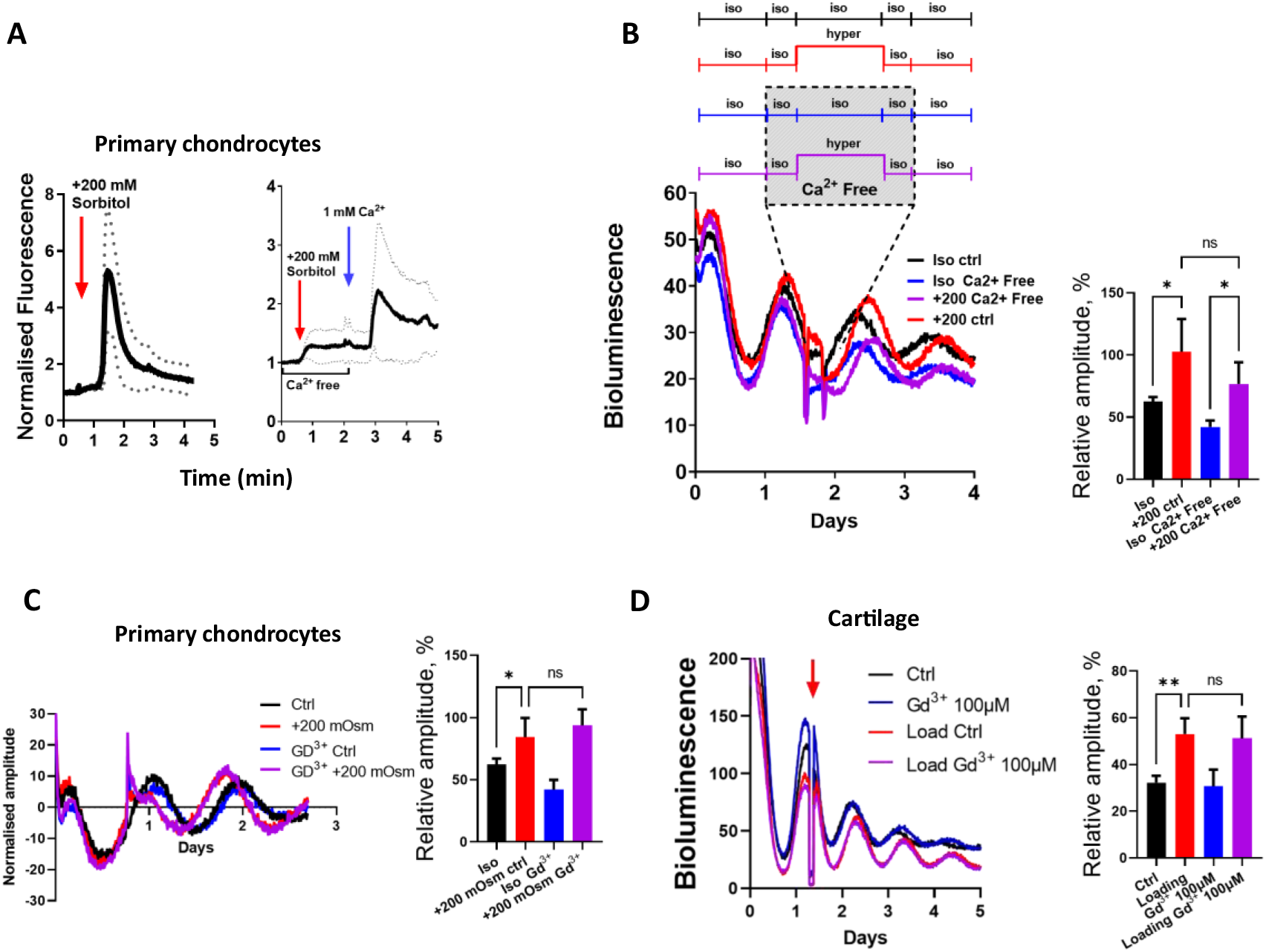
Extracellular calcium and membrane calcium channels are not involved in hyperosmolarity-induced clock resetting. (**A**) Calcium imaging in mouse primary chondrocytes using Fluo-4 fluorescence microscopy. Chondrocytes responded to +200 mOsm increase in osmolarity (red arrow) by a surge of intracellular calcium levels in calcium containing medium (left), but not in calcium free medium (right). Blue arrow indicates re-introduction of calcium to the medium, which restored calcium response. Mean ±SD of average fluorescence signals from 15 cells. (**B**) At mid-descending phase of PER2::Luc, IVD explants were pre-incubated in iso-osmotic calcium free medium (blue and purple traces). Explants were exposed to +200 mOsm increase in osmolarity for 6 hr (red and purple) and returned to iso-osmotic for 1 hr post-incubation (blue and purple traces). Subsequently media for all explants was changed to calcium containing iso-osmotic medium. Black trace-calcium containing iso-osmotic control. Each trace represents means of 4 explants. (**C**) Effect of a calcium channel blocker gadolinium on hyperosmolarity (+200 mOsm) resetting of clocks in PER2::Luc primary mouse chondrocytes. (**D**) Effect of gadolinium on loading (1 Hz, 1 hr, 0.5 MPa) induced resetting of clocks in PER2::Luc IVDs. Each trace represents mean of 3 explants. Amplitude of the peak after treatment is expressed as % of the amplitude of the peak before treatment. Mean ±SD. One-way ANOVA. *p<0.05; **p<0.01; ns- not significant.

**Figure S9.**
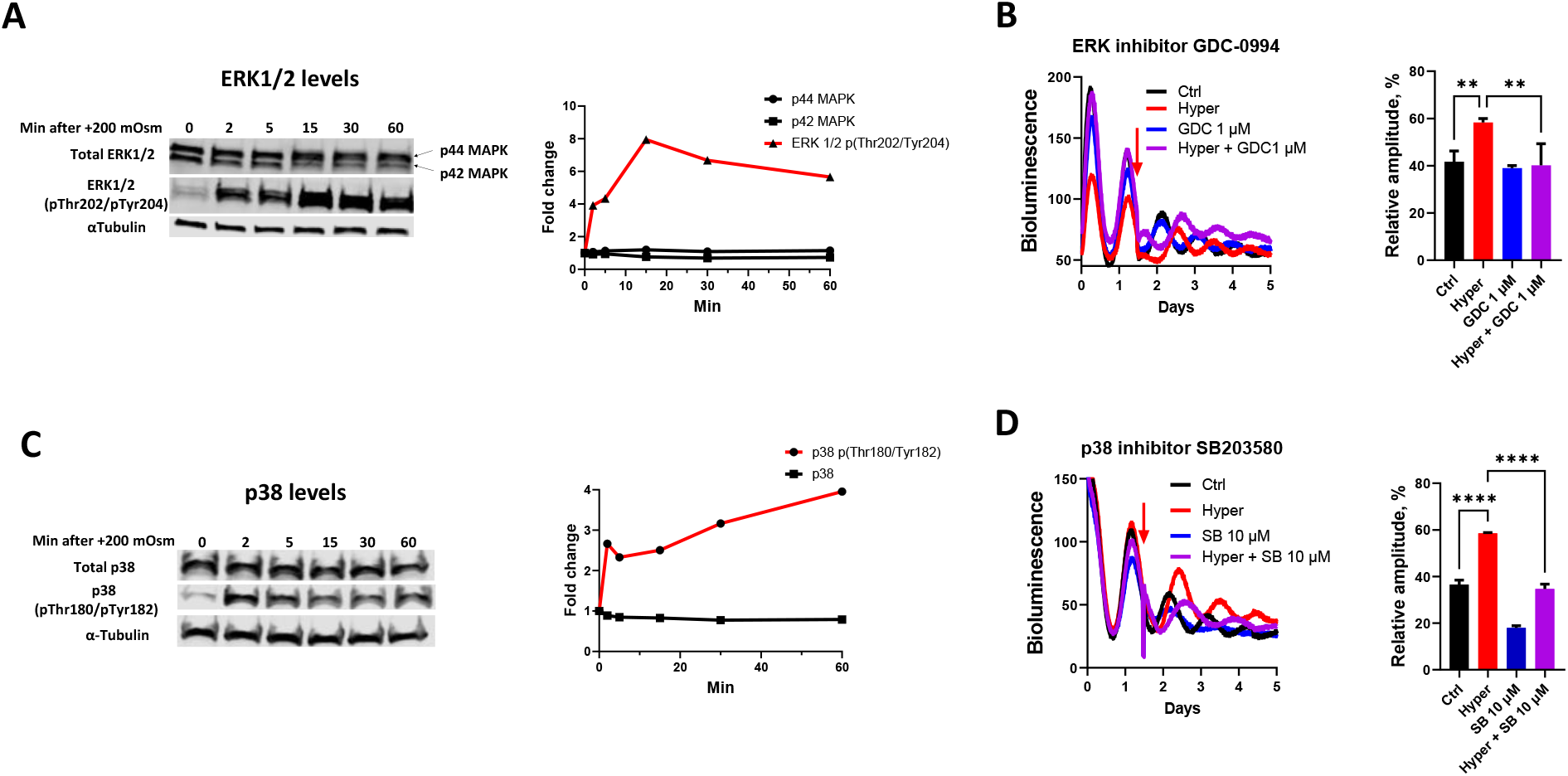
Hyperosmolarity induced activation of ERK1/2 and p38 regulate the circadian clock. **(A)** WB and quantification (relative to αTubulin) showing phosphorylation of ERK1/2 at Thr202/Tyr204 in mouse primary chondrocytes following +200 mOsm change in osmolarity. **(B)** Bioluminescence recording and amplitude quantification of PER2::Luc IVD explants. At 36 hr (mid-descending phase) explants were exposed to +200 mOsm increase in osmolarity with or without the ERK1/2 inhibitor GDC-0994. **(C)**WB and quantification (relative to αTubulin) showing phosphorylation of p38 at Thr180/Tyr182 in mouse primary chondrocytes following +200 mOsm change in osmolarity. **(D)** Bioluminescence recording and amplitude quantification of PER2::Luc IVD explants. At 36 hr (mid-descending phase) explants were exposed to +200 mOsm increase in osmolarity with or without the p38 inhibitor SB203580. Amplitude of the peak after treatment is expressed as % of the amplitude of the peak before treatment. Mean + SD. One-way ANOVA. **p<0.01; ****p<0.0001; ns- not significant.

**Figure S10.**
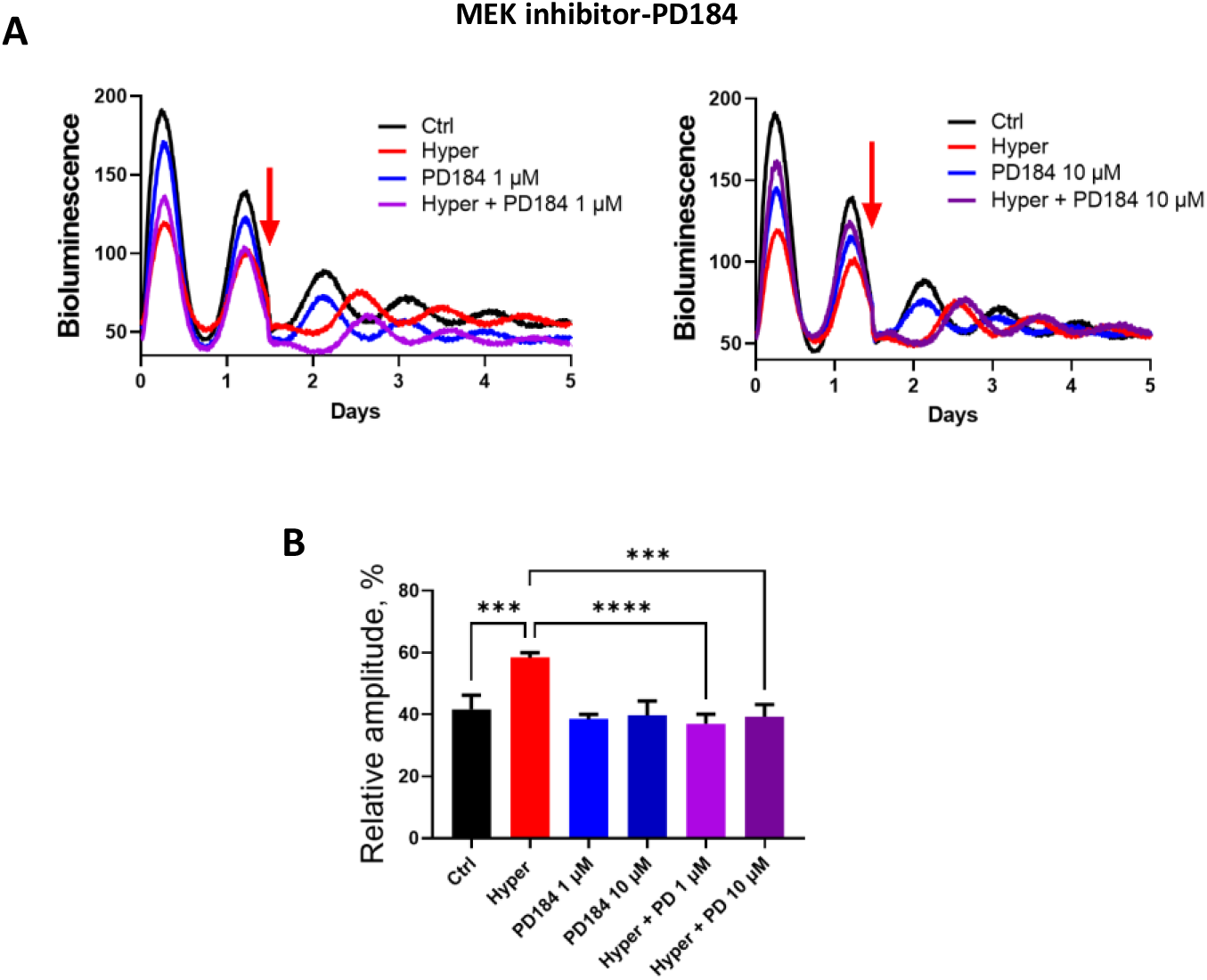
MEK inhibitor blocks the clock resetting effect of hyperosmolarity in IVDs. **(A)** PER2::Luc IVD explants were exposed to +200 mOsm increase in osmolarity (red arrow) with or without the MEK inhibitor PD184352 (purple and red trace). Each trace represents mean of 3 explants. **(B)** Amplitude after hyperosmotic challenge was quantified as % of the amplitude of the peak before challenge. Mean ± SD. One-way ANOVA. ***, P<0.001.

**Figure S11.**
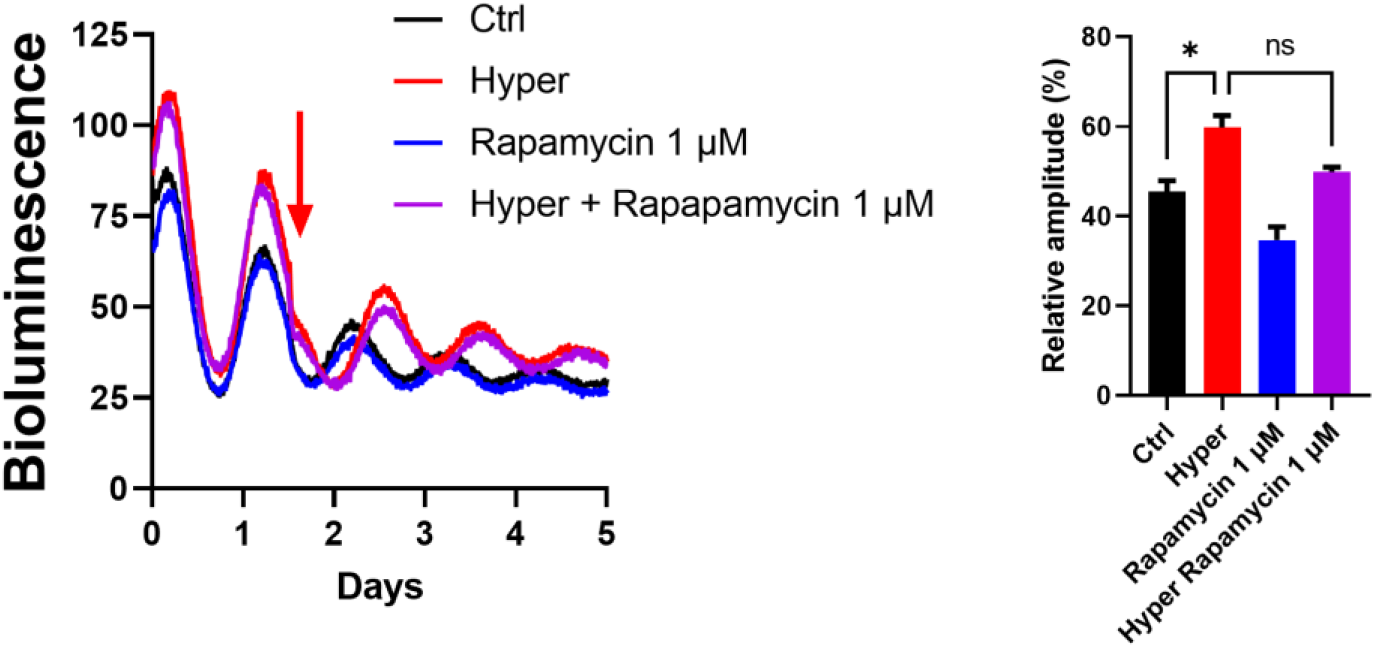
No effect of Rapamycin on hyperosmolarity induced clock resetting. At mid-descending phase PER2::Luc IVD explants were exposed to +200 mOsm increase in osmolarity (red arrow), with or without mTORC1 inhibitor rapamycin (purple and red trace). Each trace represents mean of 3 explants. Amplitude after treatment was quantified as % of the amplitude of the peak before treatment. Mean ± SD. One-way ANOVA. *p<0.05; **p<0.01; ***p<0.001; ns-not significant.

**Figure S12.**
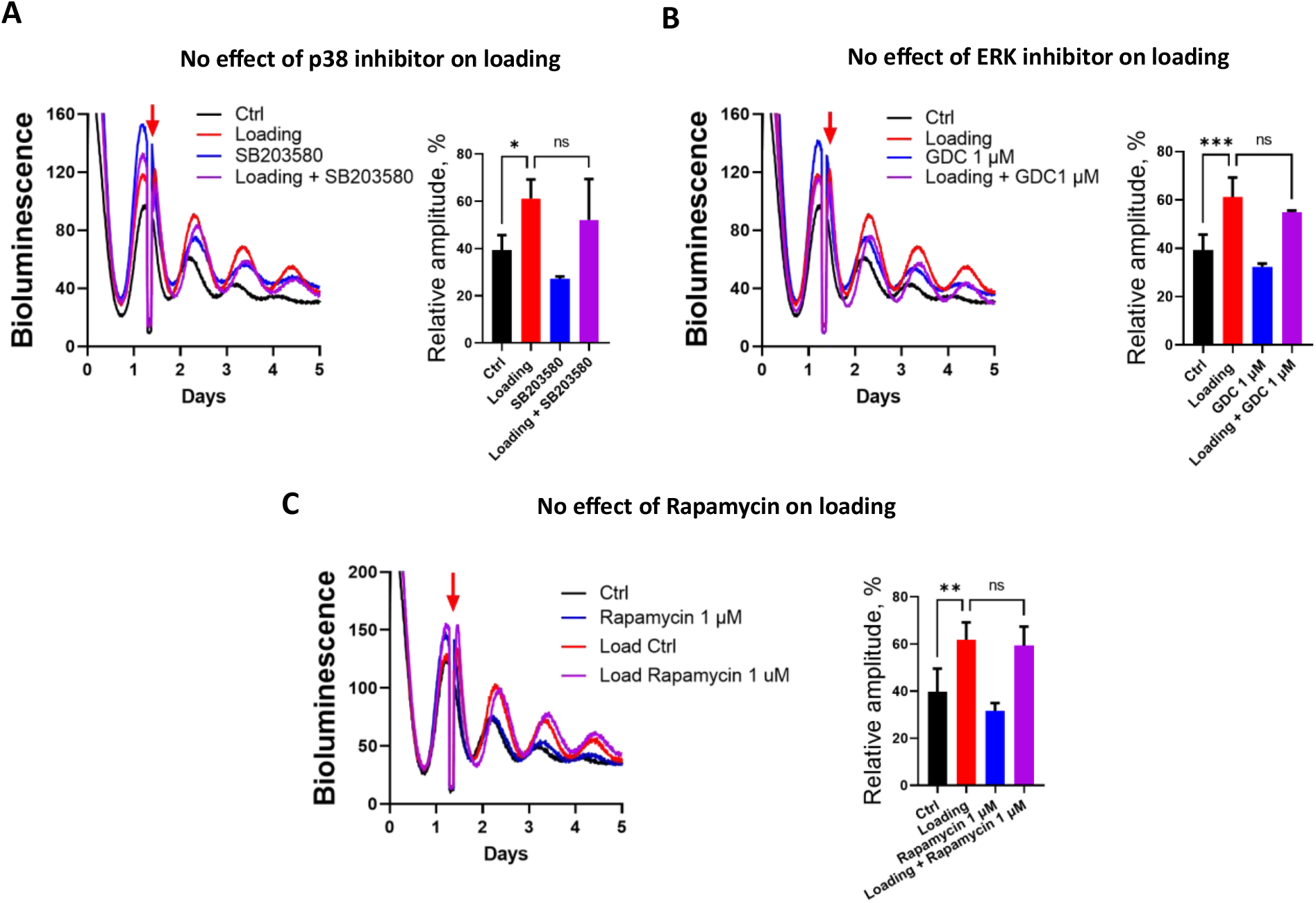
Lack of effect of a p38 inhibitor, an ERK inhibitor or Rapamycin on loading induced clock resetting in cartilage explants. Bioluminescence recording and amplitude quantification of PER2::Luc femoral head cartilage explants exposed to mechanical loading (0.5 MPa, 1 Hz, 1 hr), with or without the p38 inhibitor SB203580 (A), the ERK1/2 inhibitor GDC-0994 (B), or the mTORC1 inhibitor rapamycin (C). Each trace represents mean of 3 explants. Amplitude after loading was quantified as % of the amplitude of the peak before loading. Mean ± SD. One-way ANOVA. *p<0.05; **p<0.01; ***p<0.001; ns- not significant.

**Movie S1.**

Live bioluminescence imaging of a fragment of lumbar spine isolated from a PER2::Luc mouse using high sensitivity EMCCD camera. Bioluminescence signals in the Dexamethasone-synchronised IVDs gradually dampen in culture. The hyperosmolarity challenge (+200 mOsm) at 66 hours elicited a clear synchronization of circadian rhythms.

**Data S1. (separate file)**

List of hyperosmolarity induced rhythmic genes in primary mouse chondrocytes. pAdj<0.05.

**Data S2. (separate file)**

List of hyperosmolarity induced rhythmic genes in primary mouse chondrocytes. BHQ<0.05.

**Data S3. (separate file)**

List of hyperosmolarity induced rhythmic pathways in primary mouse chondrocytes. pAdj<0.05.

**Data S4. (separate file)**

List of differentially expressed genes between T0 and T4 after hyperosmolarity in primary mouse chondrocytes.

**Data S5. (separate file)**

Upstream regulator analysis based on the list of differentially expressed genes between T0 and T4 after hyperosmolarity in primary mouse chondrocytes.

